# Low relative air humidity leads to smaller, denser stomata and higher stomatal ratios in Arabidopsis

**DOI:** 10.1101/2025.01.31.635895

**Authors:** Ingmar Tulva, Pirko Jalakas, Elena Ivandi, Alexander Wierzchula, Patricia Kika Obinwanne, Hanna Hõrak

**Affiliations:** Institute of Technology, University of Tartu, Nooruse 1, 50411 Tartu, Estonia

**Keywords:** stomatal development, stomatal ratio, stomatal conductance, adaxial and abaxial stomata, Arabidopsis, low humidity, VPD, plant growth

## Abstract

Atmospheric dryness is increasing, bringing about decreases in plant productivity. Stomatal pores mediate plant gas-exchange with the environment, balancing CO_2_ uptake with water loss. Stomatal anatomical and physiological traits respond to changes in environment, potentially affecting plant growth and yield under future environments. Designing stomatal patterns to suit future climate conditions has the potential to improve plant water use efficiency or productivity. By combining mutations in signaling pathways that control stomatal development and apertures, we designed plants that combine high stomatal densities with more open stomata and show that respective mutations independently affect stomatal conductance and density. Analyses of adaxial and abaxial stomatal conductances showed that in Arabidopsis, adaxial stomata are responsible for a significant proportion of leaf gas-exchange. Adaxial and abaxial stomatal physiology were largely similarly affected by mutations in stomatal regulation pathways but adaxial stomata tended to be relatively more closed than abaxial stomata. We show that growth under low relative air humidity leads to higher stomatal densities and smaller stomata. Stomatal development in the adaxial and abaxial leaf surface responded differently to dry air conditions: adaxial stomatal index increased, whereas abaxial stomatal index decreased. Stomatal ratio increased under dry air conditions, leading to a higher degree of amphistomaty. Plant growth was independently suppressed by dry air and high stomatal density and index. Our results suggest that acclimation to decreasing air humidity leads to stomatal anatomical adjustments that help to maximize plant gas-exchange potential under conditions where water supply may be limited and sporadic.

## Introduction

Stomata are adjustable pores in leaves that control plant CO_2_ uptake and water loss. Stomatal density (SD), size, and aperture all impact on plant gas-exchange, with density and aperture having the largest effects on operational stomatal conductance (Ochoa et al., 2024). High stomatal densities and apertures have been linked with improved photosynthesis, particularly under fluctuating light conditions, but are accompanied by increased water loss (Tanaka et al., 2013; Wang et al., 2014; Kimura et al., 2020; Sakoda et al., 2020). Low stomatal densities and apertures lead to increased water use efficiency and/or drought tolerance in many plant species (Hepworth et al., 2015; Yang et al., 2016; Hughes et al., 2017; Caine et al., 2019; Dunn et al., 2019; Mega et al., 2019; Xiao et al., 2024).

Atmospheric dryness has been rising throughout and is expected to increase further in the current century, leading to yield losses (Yuan et al., 2019; López et al., 2021). Learning how plant stomatal traits respond to dry air conditions is important to understand plant behavior in future climates. Drier air means higher vapor pressure deficit (VPD, the difference between real and saturating air humidity), which triggers stomatal closure in plants (Jalakas et al., 2021). The effects of VPD on stomatal density are less clear. In several plant species SD increased under high relative air humidity (RH) conditions (Bakker, 1991; Torre et al., 2003), whereas further studies indicated that SD response to VPD is species-specific (reviewed in Fanourakis et al., 2020). In Arabidopsis, growth under high RH resulted in lower SD (Arve et al., 2015), whereas dry air conditions increased SD (Tulva et al., 2024).

In addition to stomatal density, aperture, and size, in amphistomatous plants such as Arabidopsis, stomatal distribution between adaxial (upper) and abaxial (lower) leaf surfaces can also affect gas exchange. Stomatal distribution between leaf surfaces is characterised by stomatal ratio (SD ratio, ratio of adaxial and abaxial SD), which depends on plant genotype and environment. In some plant species, adaxial and abaxial stomatal anatomical traits respond differently to growth at high RH (Fanourakis et al., 2020), suggesting a different developmental response to environmental conditions in the upper and lower leaf surface. In line with this, high light and dry air increase stomatal ratio (Hronková et al., 2015; Devi and Reddy, 2018; Tulva et al., 2024). How the adjustment of stomatal ratio in response to environmental cues occurs in plants and what is its functional role, remains largely unknown.

Stomatal development in plants is regulated by a concerted action of transcription factors, signaling peptides, their receptors, and various kinases (Zoulias et al., 2018). The epidermal patterning factors EPF1 and EPF2 are peptides that ensure proper stomatal spacing in the epidermis by preventing adjacent stomatal formation (Hara et al., 2007; Hunt and Gray, 2009). Lack of both of these peptides leads to increased SD in Arabidopsis and the production of arrested stomatal precursor cells in the abaxial epidermis (Hunt and Gray, 2009; Jalakas et al., 2024).

Stomatal apertures depend on environmental conditions and are controlled by guard cell signaling pathways that trigger stomatal opening or closure. Major stomatal closure signaling pathways include the abscisic acid (ABA) and CO_2_ signaling pathways (Dubeaux et al., 2021; Hsu et al., 2021a). ABA is induced in response to drought and dry air conditions (McAdam et al., 2016; Kuromori et al., 2022) and triggers stomatal closure via a pathway mediated by ABA receptors, protein phosphatases, and the kinase OPEN STOMATA 1 (OST1) that phosphorylates the major guard cell slow-type anion channel SLOW ANION CHANNEL-ASSOCIATED 1 (SLAC1), leading to ion efflux from guard cells and stomatal closure (Negi et al., 2008; Vahisalu et al., 2008; Fujii et al., 2009; Geiger et al., 2009; Ma et al., 2009; Park et al., 2009). The pseudokinase GUARD CELL HYDROGEN PEROXIDE-RESISTANT 1 (GHR1) also contributes to ion channel activation in this pathway, potentially as a scaffold that brings together proteins that activate SLAC1 (Hua et al., 2012; Sierla et al., 2018). The ABA signaling pathway largely mediates stomatal response to elevated VPD and is involved in responses to other environmental cues (Merilo et al., 2013). In a CO_2_-specific branch of stomatal closure pathway, the kinases HIGH LEAF TEMPERATURE 1 (HT1) and MITOGEN-ACTIVATED PROTEIN KINASE 12 (MPK12) form the sensor that perceives changes in CO_2_ levels and regulates SLAC1 activation (Hashimoto et al., 2006; Hõrak et al., 2016; Jakobson et al., 2016; Takahashi et al., 2022). Loss of MPK12, GHR1, OST1, or SLAC1 function leads to increased stomatal conductance due to more open stomata (Vahisalu et al., 2008; Merilo et al., 2013; Jakobson et al., 2016; Sierla et al., 2018). By combining alleles that lead to large stomatal apertures with those that lead to high SDs, it may be possible to increase plant photosynthetic capacity and biomass production.

Here, we generated plant lines that combine high SD with very open stomata, and analysed the effect of respective traits on plant gas-exchange, epidermal patterning, and growth. We found independent effects of mutations that affect stomatal development and physiology on stomatal density and conductance. Adaxial and abaxial stomatal development were differently regulated in Arabidopsis, whereas leaf-side specific stomatal conductance largely reflected the distribution of stomata between adaxial and abaxial leaf surfaces. We show that low relative air humidity stimulates adaxial stomatal production and increases stomatal ratio, while both low humidity and high SD independently suppress plant growth.

## Results

### Mutations affecting stomatal density and openness act independently

We aimed to test if signaling pathways that control plant stomatal development and function act independently by combining mutations known to affect either process in Arabidopsis. To this end, we generated mutants expected to have increased stomatal densities due to the lack of function of both EPF1 and EPF2, and very open stomata due to deficiency of MPK12, GHR1, OST1, or SLAC1. We analysed SD of the obtained triple mutants, and found that the mutant *EPF1* and *EPF2* alleles increased both adaxial and abaxial SD similarly in the background of all other studied genotypes (Figure 1A-B). Whole-rosette leaf conductance (g_l_) was significantly increased in the *epf1/2* background by all mutations that also increased g_l_ in the wild-type background (*mpk12-4*, *ghr1-3*, *ost1-3*, Figure 1C), indicating independent effects of impairments in signaling pathways that control stomatal development and aperture regulation on plant stomatal density and conductance. Net assimilation rate (A_net_) was largely similar between all studied plant lines, with a notable decrease in A_net_ in the *ghr1-3epf1/2* mutant compared with the *epf1/2* mutant (Figure 1D). The ratio of adaxial to abaxial SDs (SD ratio) was overall decreased by the lack of EPFs (significant main effect in Table 1, Figure 1E). Total SD (sum of adaxial and abaxial SDs) was positively related with g_l_ (Figure 1F), whereas there was no significant relationship between g_l_ and A_net_ or SD and A_net_ (Supplementary Figure S1).

**Figure 1.**
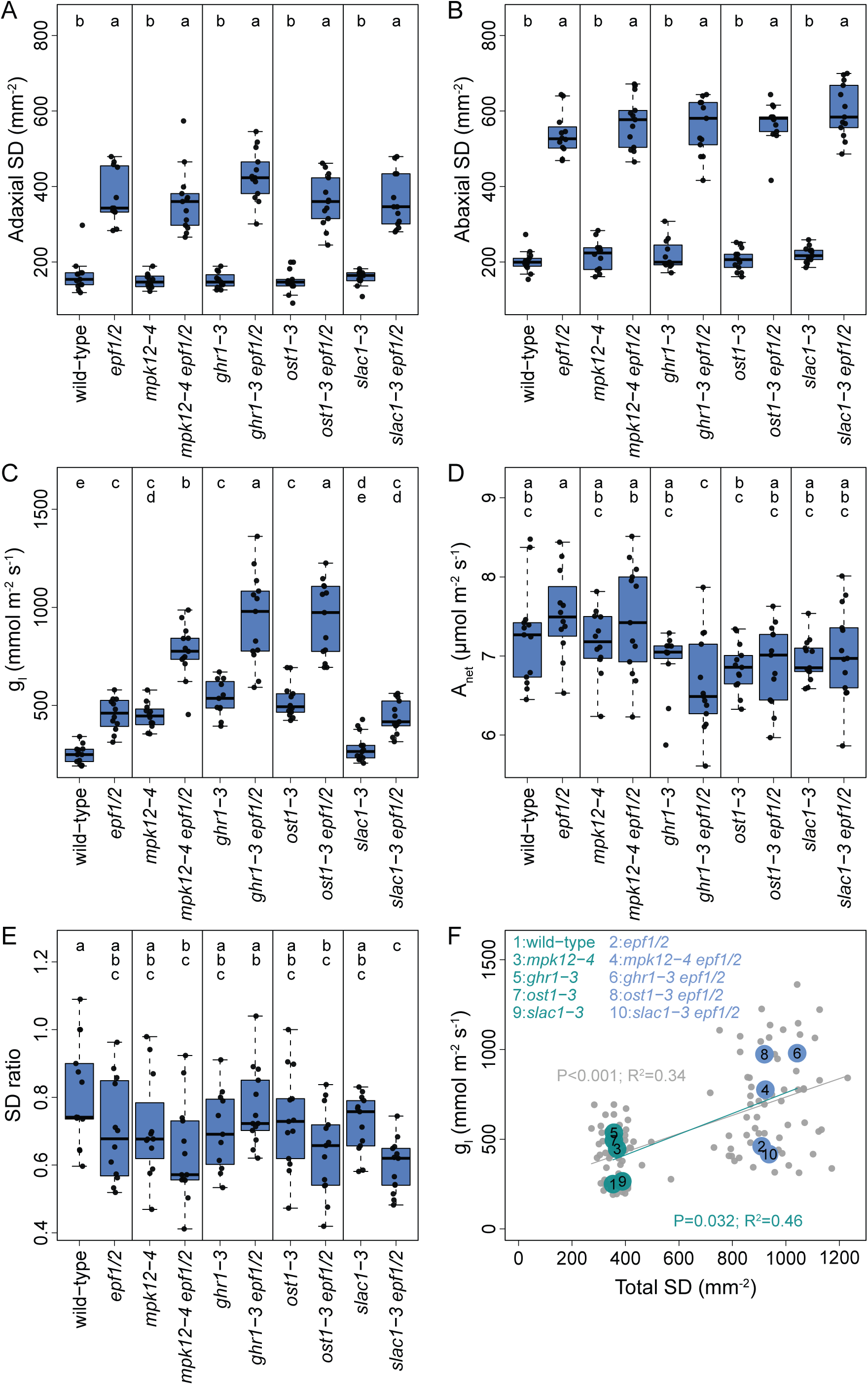
Plant stomatal anatomy and physiology in the gas exchange experiment. (A)-(B), stomatal density of each leaf side; (C) leaf conductance (i.e. stomatal plus cuticular conductance); (D), net photosynthesis; (E), stomatal density ratio. For each plant line (n=11-13), the box denotes the upper to lower quartile range, with the median marked with a line. The whiskers denote the non-outlier range, and black dots mark individual data points. Letters above the boxes denote Tukey post-hoc test results, with plant lines not sharing a letter differing from each other at p<0.05. (F), relationship between total SD (sum of both leaf sides) and leaf conductance. Grey points are individual plants, large numbered dots in the foreground show genotype medians.

**Table 1.**
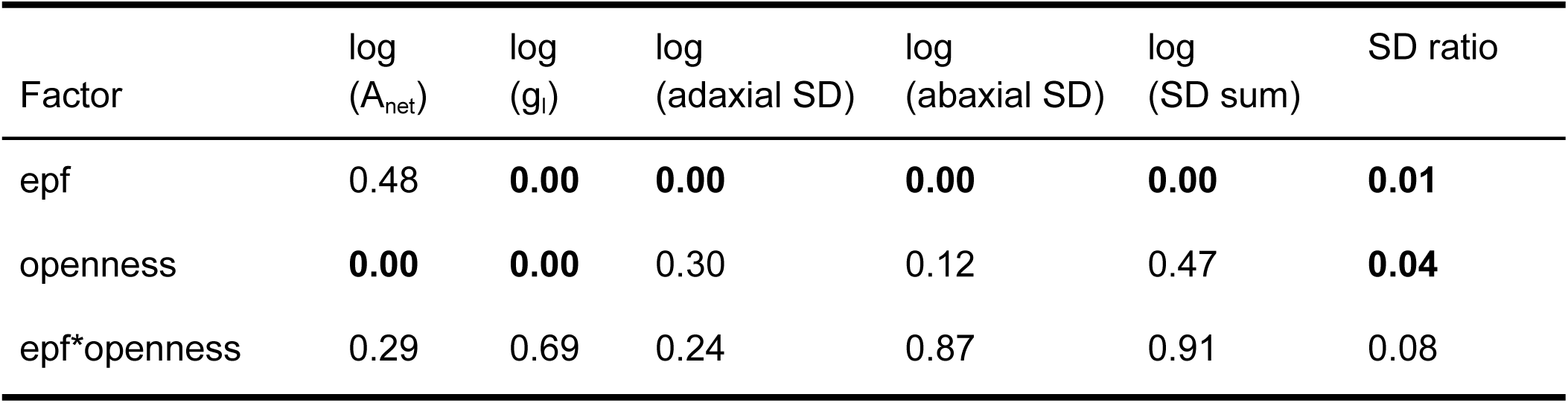
Results of analysis of variance for whole-rosette gas exchange. Shown are p-values of respective factors. A_net_, net CO_2_ assimilation; g_l_, leaf conductance (sum of stomatal and cuticular conductance); SD, stomatal density; SD ratio, ratio of upper to lower SD. Log-transformed values were analysed when the raw variable failed Levene’s test for homogeneity of variance.

| Factor | log<br>($A_{\text{net}}$) | log<br>($g_l$) | log<br>(adaxial SD) | log<br>(abaxial SD) | log<br>(SD sum) | SD ratio |
| --- | --- | --- | --- | --- | --- | --- |
| epf | 0.48 | **0.00** | **0.00** | **0.00** | **0.00** | **0.01** |
| openness | **0.00** | **0.00** | 0.30 | 0.12 | 0.47 | **0.04** |
| epf*openness | 0.29 | 0.69 | 0.24 | 0.87 | 0.91 | 0.08 |

### Adaxial and abaxial stomatal conductances follow stomatal distribution between leaf surfaces

To independently address adaxial and abaxial stomatal function in our studied mutants, we measured their adaxial and abaxial stomatal conductance (g_s_) with a porometer. Adaxial g_s_ accounted for a substantial proportion of total leaf conductance (∼40% in wild-type plants, Supplementary Figure S2), and was increased by the *epf1/2* mutations and by mutations affecting stomatal apertures (significant ‘epf’ and ‘openness’ main effects in Table 2, Figure 2A). Abaxial g_s_ was increased by the *epf1/2* mutations in all backgrounds except wild-type (Figure 2B), and overall increased by mutations that affected stomatal openness (significant ‘openness’ main effect, Table 2). Leaves that were used for porometry showed similarly high SD values for all mutants harboring the *epf1/2* alleles and no effect of other mutant alleles on SD (Supplementary Figure S2). Both SD ratios and stomatal conductance ratios (g_s_ ratio, ratio of adaxial and abaxial g_s_) were decreased by the *epf1/2* alleles (significant main effects in Table 2, Figure 2C-D), whereas g_s_ ratio was also affected by stomatal openness (significant epf*openness interaction in Table 2, Figure 2D). Across genotypes, there was a significant positive relationship between SD ratio and g_s_ ratio (Figure 2E).

**Figure 2.**
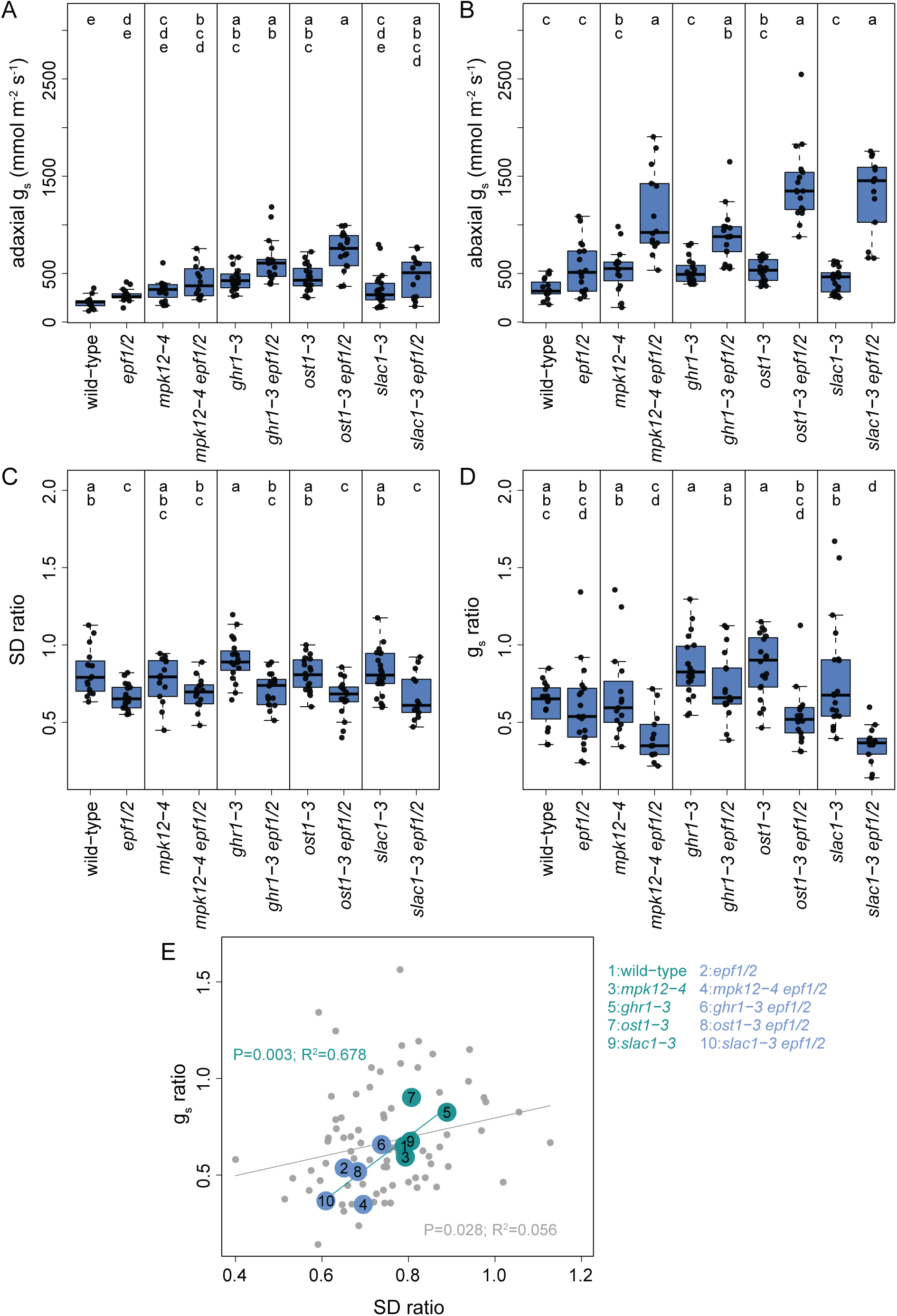
Stomatal behaviour in the porometry experiment. (A)-(B), stomatal conductance of a healthy leaf; adaxial to abaxial side ratio of stomatal densities (C) and conductances (D). For each plant line (n=14-22), the box denotes the lower to upper quartile range, with the median marked with a line. The whiskers span the non-outlier range, and black dots mark individual data points. Letters above the boxes denote Dunn’s (panels A, B) or Tukey’s (panels C, D) post-hoc test results, with plant lines not sharing a letter differing from each other at p<0.05. (E), relationship between SD ratio and g_s_ ratio. Grey dots denote individual data points from one batch (in the other, individual correspondence between SD and g_s_ was lost in sample handling); large numbered circles mark genotype medians from two batches pooled.

**Table 2.**
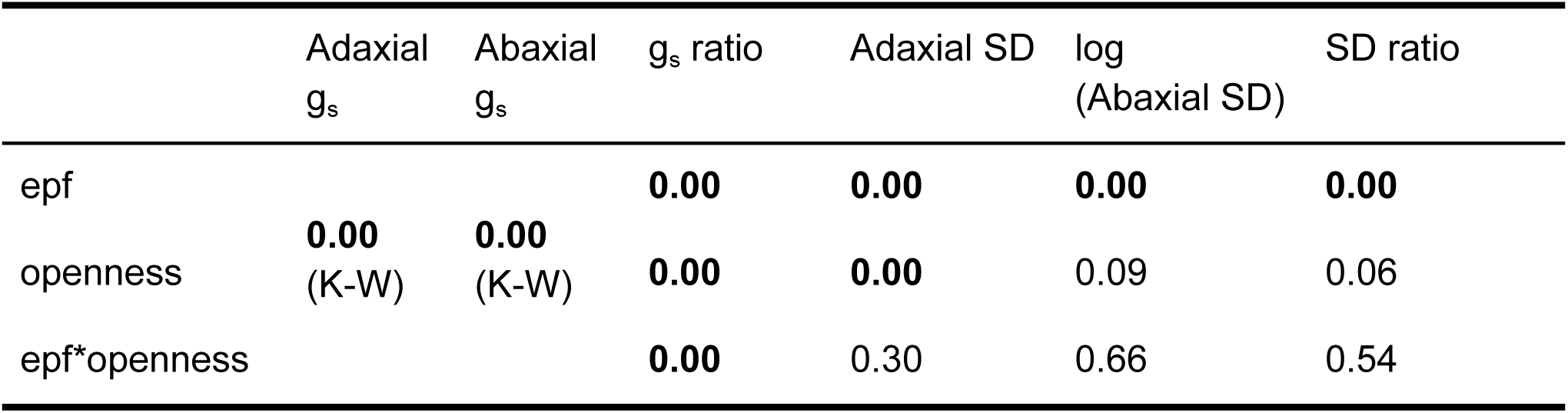
Results of analysis of variance for the porometry experiment. Shown are p-values of respective factors on each variable in the title row. g_s_, stomatal conductance as shown by the LI-600 porometer; SD, stomatal density; g_s_ ratio is the ratio of adaxial to abaxial conductance, and SD ratio is the ratio of adaxial to abaxial SD. Log-transformed values were analysed with ANOVA where the raw variable failed Levene’s test for homogeneity of variance. In the case of stomatal conductances (g_s_), the log-transformed variables still failed Levene’s test; for them, Kruskal-Wallis was performed with genotype as the predictor, and the results of this analysis are shown.

### Growth under dry air conditions leads to smaller denser stomata

We next aimed to test how plants with increased stomatal densities and conductances develop and grow under normal or low relative air humidity (RH) conditions, and how low RH affects stomatal anatomical traits. Growth under low RH led to increased SDs on both adaxial and abaxial leaf surfaces (Tables 3 and 4, Figure 3A-B), whereas stomata were smaller under dry air conditions on both leaf surfaces (Table 3, Figure 3C-D). Under control conditions, adaxial stomata were larger in plants carrying the *ost1-3*, *slac1-3*, and *mpk12-4* alleles, whereas under low RH, plants carrying the *ghr1-3* allele had the smallest adaxial stomata (significant RH*sensitivity effect in Table 3, Supplementary Table S3, Figure 3C). No such interaction was found for the abaxial leaf surface (Table 3). There was a negative relationship between SD and stomatal size when separately addressing this in mutants with or without the *epf1/2* alleles on both leaf surfaces, whereas there was no significant association between these traits across all studied genotypes (Figure 3E-F).

**Figure 3.**
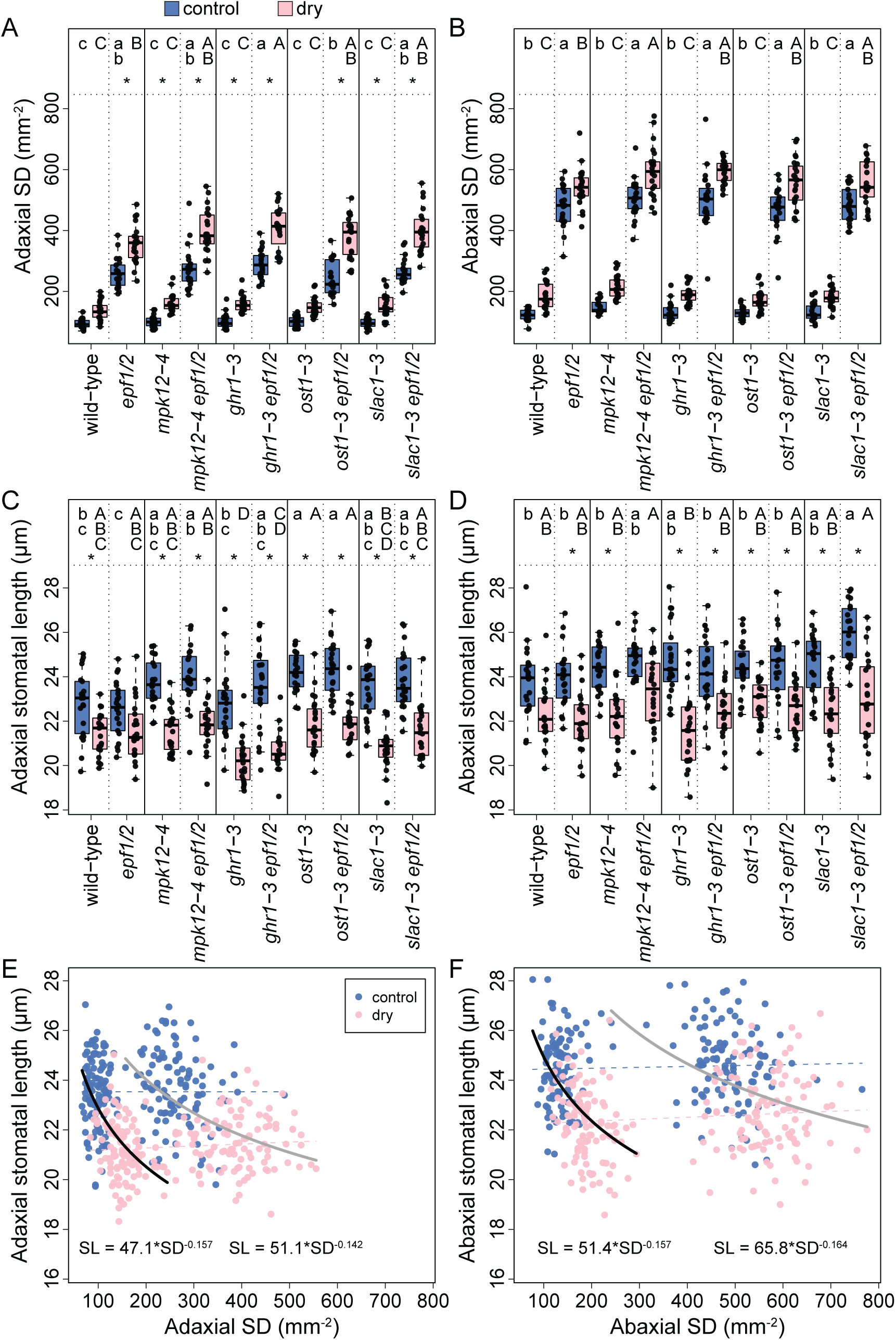
Stomatal anatomy in the growth experiment. (A)-(B), stomatal density on the adaxial and abaxial side; (C)-(D), stomatal length in the same samples as (A) and (B), respectively. The boxes denote the lower to upper quartile range, with the median marked with a line. The whiskers span the non-outlier range, and black dots mark individual data points (n=19-23). Asterisk above a pair of boxes means significant pairwise difference between plants grown in low RH and control conditions according to Tukey test (factorial ANOVA involving all treatment and line combinations). Small letters above the blue control boxes denote Tukey (Dunn for panel B) post-hoc test results, with plant lines not sharing a letter differing from each other at p<0.05 (one-way ANOVA for genotype differences involving only control plants), and capital letters analogously for the low RH treatment. (E)-(F), relationship between stomatal density and stomatal size for each leaf side. Blue dots denote control and pink dots low RH-grown plants; the two distinctive point clouds on each panel are due to *epf1/2* mutation in half the plants and dashed lines show linear regressions, whose slopes did not differ significantly from zero. The black (for plant lines with intact EPFs) and grey (for *epf1/2* mutants) denote power regressions (Table S2) where plants from different growth RH were pooled.

**Table 3.** Results of analysis of variance for the growth experiment. Shown are p-values of each factor or their interaction. SD, stomatal density; SI, stomatal index; LCI, stomatal lineage cell index. Log-transformed variables were analysed when the raw variable failed Levene’s test for homogeneity of variance.

|  | log<br>(dry mass) | LMA | log<br>(adax. SD) | SD<br>ratio | log<br>(total<br>LCD) | adaxial<br>SI | abaxial<br>SI | adaxial<br>stomatal<br>length | abaxial<br>stomatal<br>length | adaxial<br>LCI | abaxial<br>LCI |
| --- | --- | --- | --- | --- | --- | --- | --- | --- | --- | --- | --- |
| RH | **0.00** | **0.00** | **0.00** | **0.00** | **0.00** | **0.01** | **0.00** | **0.00** | **0.00** | **0.00** | 0.74 |
| epf | **0.00** | 0.90 | **0.00** | **0.00** | **0.00** | **0.00** | **0.00** | **0.02** | **0.01** | **0.00** | **0.00** |
| openness | **0.00** | **0.00** | **0.00** | **0.03** | **0.03** | **0.00** | **0.00** | **0.00** | **0.00** | **0.00** | **0.00** |
| RH*epf | 0.58 | 0.34 | 0.16 | **0.01** | 0.09 | 0.57 | **0.00** | 0.69 | 0.67 | 0.71 | 0.94 |
| RH*openness | 0.32 | 0.12 | 0.44 | 0.16 | 0.39 | 1.00 | 0.60 | **0.00** | 0.16 | 0.98 | 0.53 |
| epf*openness | 0.15 | 0.68 | 0.24 | 0.12 | 0.12 | **0.02** | 0.23 | 0.37 | 0.05 | **0.01** | 0.25 |
| RH*epf*openness | 0.56 | 0.72 | 0.57 | 0.69 | 0.54 | 0.51 | 0.51 | 0.78 | 0.10 | 0.56 | 0.21 |

**Table 4.** Results of Kruskal-Wallis tests for the growth experiment for variables where both the raw value and its logarithm failed Levene’s test for homogeneity of variance. Three analyses are shown: test for the RH main effect (with all genotypes pooled), and tests for differences between genotypes for each growth RH separately.

|  | Abaxial<br>SD | Adaxial<br>precursor<br>index | Abaxial<br>precursor<br>index |
| --- | --- | --- | --- |
| RH<br>(genotypes pooled) | **0.00** | **0.00** | **0.00** |
| genotype<br>(control treatment) | **0.00** | **0.00** | **0.00** |
| genotype<br>(dry treatment) | **0.00** | **0.00** | **0.00** |

### Dry air increases stomatal ratio and adaxial stomatal index

Both SD ratio (reflecting mature stomata) and stomatal lineage cells ratio (LCD ratio, calculated as the ratio of adaxial and abaxial stomatal lineage cell densities, LCDs) increased significantly in plants grown under low RH (significant main effect of RH in Table 3, Figure 4A, Supplementary Figure S3A). Both adaxial and abaxial stomatal index (SI) and stomatal lineage cell index (LCI) were increased by the *epf1/2* mutations (significant main effects in Table 3, Figure 4B-C, Supplementary Figure S3B-C). Adaxial SI was slightly increased by low RH and abaxial SI was decreased by low RH, whereas LCI was significantly increased only in the adaxial leaf surface (significant main effects in Table 3, Figure 4B-C, Supplementary Figure S3B-C). The *ghr1-3* mutants had lower SI and LCI compared with wild-type in the adaxial epidermis (Figure 4B, Supplementary Figure S3B).

**Figure 4.**
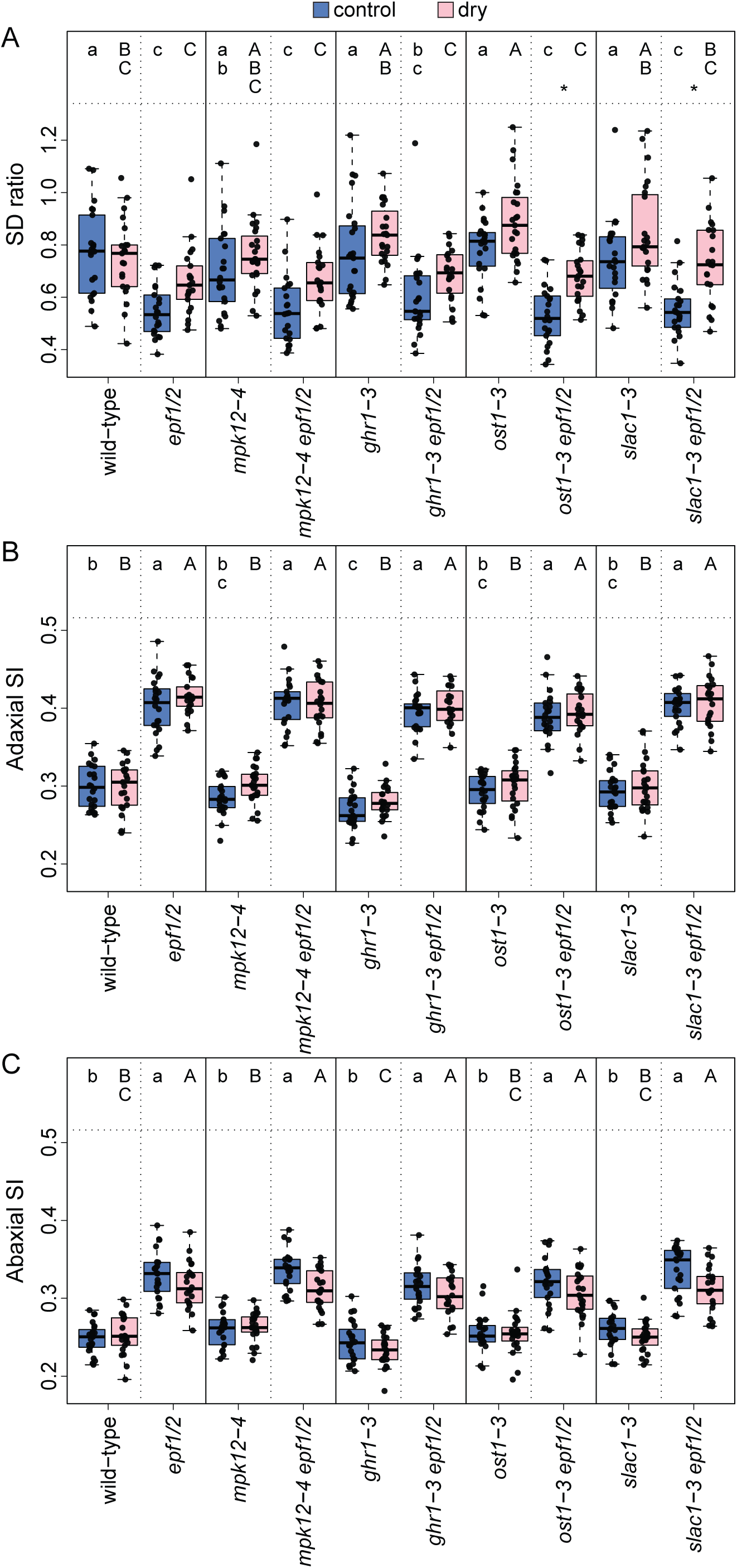
Stomatal density ratio between adaxial and abaxial side (A) and stomatal indices on both leaf sides (B)-(C). The boxes denote the lower to upper quartile range, with the median marked with a line. The whiskers span the non-outlier range, and black dots mark individual data points (n=20-23). Asterisk above a pair of boxes means significant pairwise difference between plants grown in low RH and control conditions according to Tukey test (factorial ANOVA involving all treatment and line combinations). Small letters above the blue control boxes denote Tukey post-hoc test results, with plant lines not sharing any letter differing from each other at p<0.05 (one-way ANOVA for genotype differences involving only control plants), and capital letters analogously for the low RH treatment.

### Stomata arrest in precursor state more often under dry air conditions

Under control conditions, arrested stomatal precursors were typically detected only in the abaxial epidermis of plants with the *epf1/2* mutations (Figure 5, Supplementary Figure S4). Under low RH, arrested stomatal precursors were found more often in both the adaxial and abaxial epidermis, although adaxial stomatal precursors still remained rare (Figure 5).

**Figure 5.**
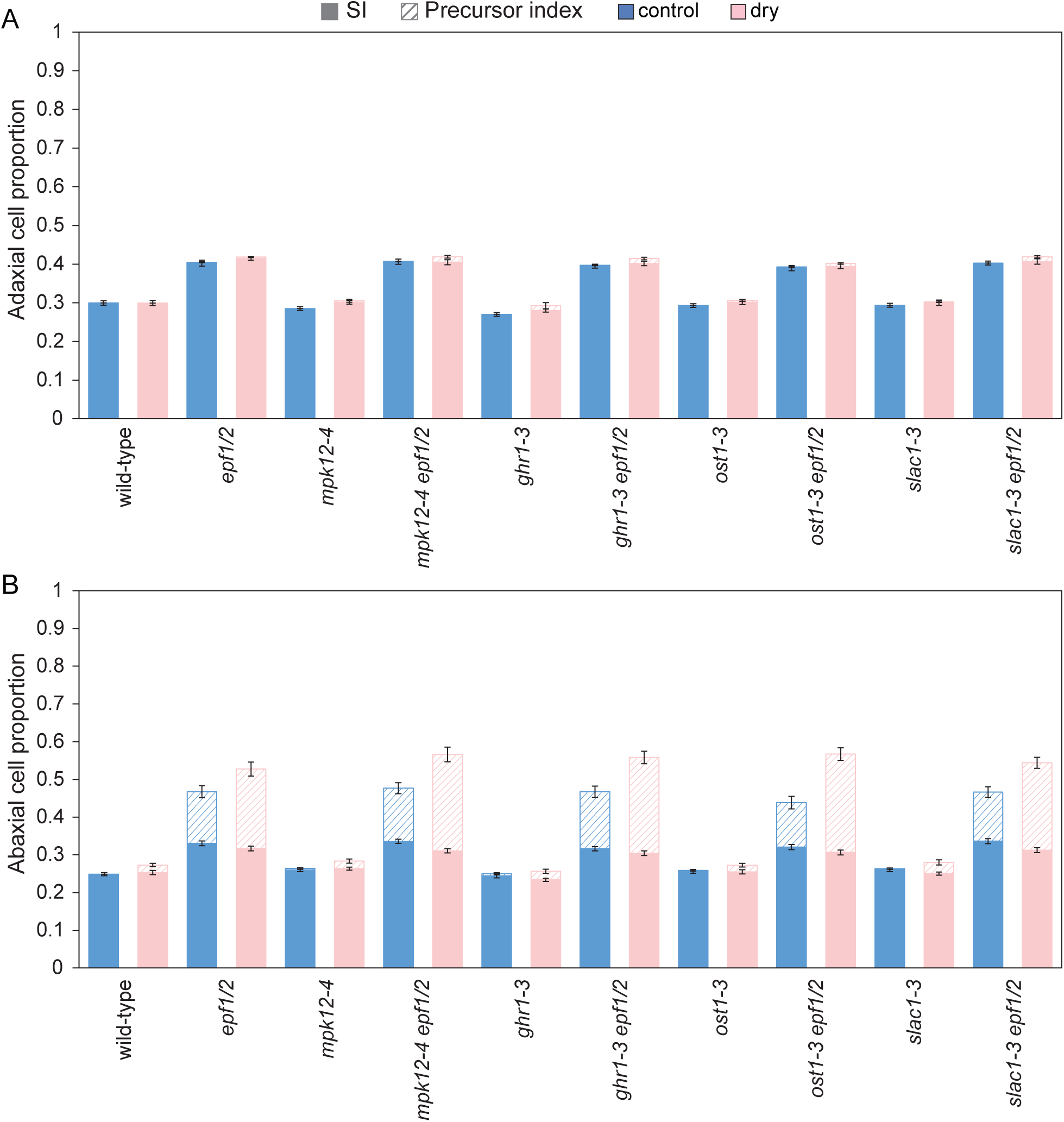
Abundance of mature stomata (i.e. SI, filled bars) and stomatal precursors (i.e. precursor index, hatched bars) among epidermal cells. Means±SE are shown (n=20-23).

### High stomatal density and index are negatively associated with plant growth

Leaf mass per area (LMA) was higher under low RH and in the *ost1-3*, *ghr1-3*, and *mpk12-4* mutants (Supplementary Figure S5, Supplementary Table S4). Plant dry mass was lower in plants grown under low RH conditions and in plants deficient in EPF1/2 function, whereas low RH and high stomatal density independently suppressed growth (significant main effects in Table 3, Figure 6A). Both SD and stomatal lineage cell index (LCI) were negatively related with growth under both growth regimens (Figure 6B-C).

**Figure 6.**
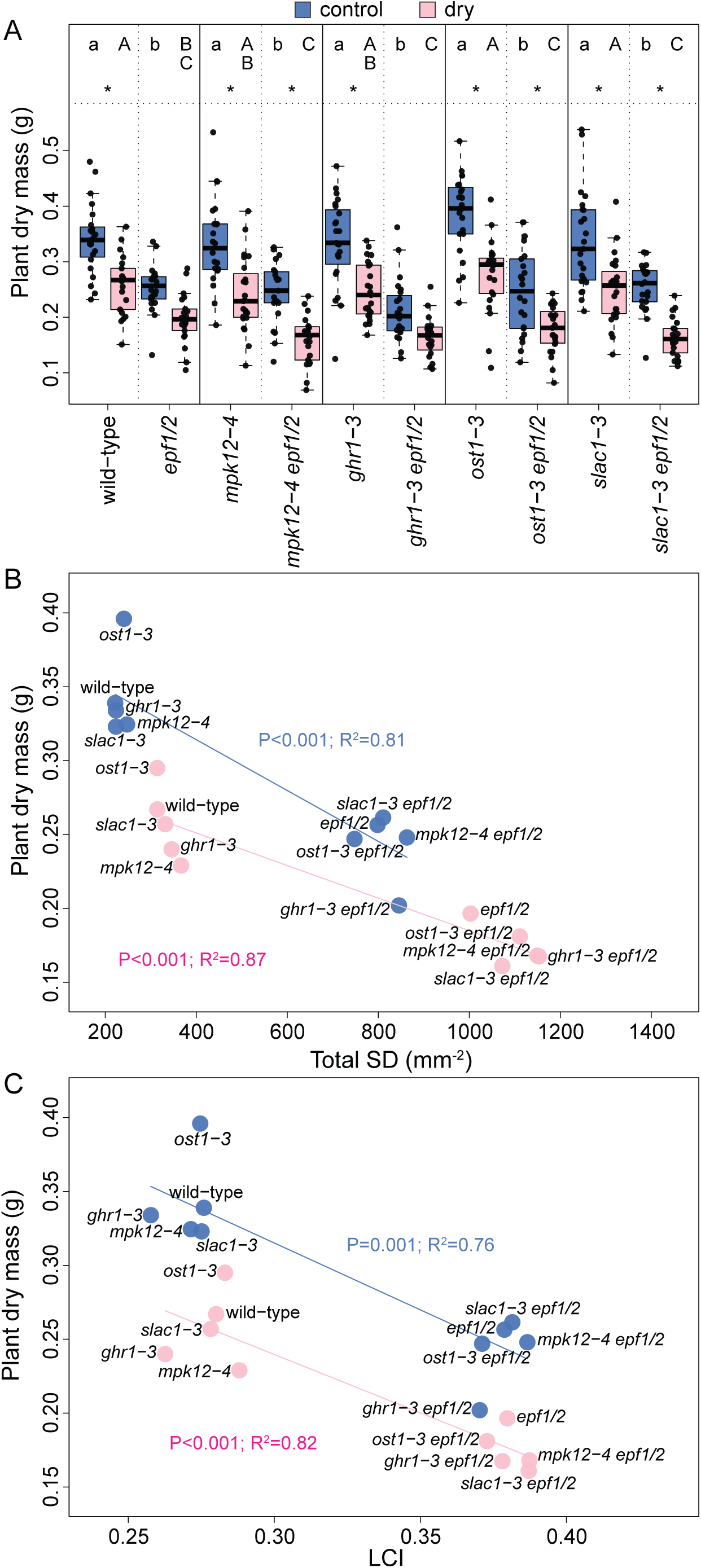
Plant above-ground dry mass (A) and its relationship with mature stomatal density (B) as well as stomatal lineage cell index (C). In (A), the boxes denote the lower to upper quartile range, with the median marked with a line. The whiskers span the non-outlier range, and black dots mark individual data points (n=20-23). Asterisk above a pair of boxes means significant pairwise difference between plants grown in low RH and control conditions according to Tukey test (factorial ANOVA involving all treatment and line combinations). Small letters above the blue control boxes denote Tukey post-hoc test results, with plant lines not sharing any letter differing from each other at p<0.05 (one-way ANOVA for genotype differences involving only control plants), and capital letters analogously for the low RH treatment. In (B) and (C), the dots denote genotype medians for given growth RH.

## Discussion

By combining mutations expected to increase both stomatal openness and density, we found that respective signaling pathways act largely independently, as the obtained mutants showed expected increases in both stomatal conductance and SD (Figures 1-2). This matches our previous finding, where a combination of mutations leading to high SDs or to more closed stomata also showed largely independent effects on both stomatal conductance and density (Tulva et al., 2024). Thus, combining mutations that affect either stomatal development or closure/opening pathways may be a viable strategy to achieve desired trait combinations, in line with previously suggested manipulations of multiple plant traits to improve photosynthesis and/or water use efficiency (Flexas, 2016). However, our studies show that increasing stomatal densities to increase CO_2_ uptake capacity at least to the degree achieved by the *EPF1/2* double loss-of-function is not a useful strategy for improving plant biomass production, as both SD and SI were negatively associated with plant growth here (Figure 6) and SD was negatively related with growth also in our previous studies (Jalakas et al., 2024; Tulva et al., 2024). Nevertheless, the additive and independent effects of the mutations in pathways studied here suggest that a similar strategy may become useful to develop plants with increased drought tolerance by combining mutations that lead to low SDs and small stomatal apertures. In addition, the *epf1/2* alleles used by us led to rather extreme increases in SD and using alleles that lead to more moderate SD increases may still prove useful to increase plant CO_2_ uptake capacity, productivity and/or growth.

Stomatal conductance is often positively correlated with A_net_ (Wong et al., 1979), and high stomatal conductance has been associated with improved biomass growth and yield (Roche, 2015). We expected that the combination of more open stomata with high SD-s would lead to higher A_net_ in plants due to improved CO_2_ uptake, even if at the cost of increased water loss. However, A_net_ was similar despite very different stomatal traits in our studied mutants (Figure 1) and not significantly associated with g_l_ (Supplementary Figure S1), suggesting that A_net_ was saturated already in plants with wild-type stomatal traits under the light conditions used in our experiments.

Genotype effects on adaxial and abaxial stomatal conductance were similar to whole-rosette leaf conductance, whereas the total sum of adaxial and abaxial stomatal conductance as measured by porometry was larger than whole-rosette g_l_ (Figures 1-2). This difference may be explained by the use of a relatively young physiologically active leaf in the porometer experiments, whereas whole-rosette gas-exchange was analysed at the stage when first leaves with potentially lower physiological activity (Stessman et al., 2002; Pantin et al., 2012) formed a large part of the rosette and may have contributed to the observed lower overall g_l_ values in these experiments. Adaxial g_s_ accounted for ∼40% of total g_s_ in the porometer experiment (Supplementary Figure S2), underlining the importance of adaxial stomata for plant gas-exchange not only in wheat, as observed before (Wall et al., 2022), but also in Arabidopsis that has less stomata on the adaxial compared with abaxial leaf surface. Thus, further studies of adaxial stomatal development and function are needed to understand their role in plant gas-exchange.

Although adaxial and abaxial stomatal conductances generally followed stomatal distribution between leaf surfaces (Figure 2), the g_s_ ratio values were often lower than respective SD ratio values, whereas these ratios appeared similar in the *ost1-3* and *ghr1-3* single mutants impaired in the ABA-induced stomatal closure pathway (Figure 2C-D). These data suggest that adaxial stomata are normally relatively more closed than abaxial stomata, whereas adaxial stomata may remain more open in plants deficient in elevated VPD-induced stomatal closure (Merilo et al., 2013; Hsu et al., 2021b) that are insensitive to the relatively drier air at the adaxial leaf surface.

Stomatal ratios were lower in plants with *epf1/2* mutations, whereas the effect was clearer in leaf 12 (Figure 4) than in leaf 6 (Figure 1). This aligns with our previous findings, where SD ratio was reduced by the *epf1/2* mutations in leaf 8 (Jalakas et al., 2024), but not in leaf 6 (Tulva et al., 2024). Although we did not detect between-leaf differences in SD ratio in Arabidopsis leaves 6, 8, 10, and 12 before (Jalakas et al., 2024), the *epf1/2* mutant was not part of this experiment.

Taken together, our experiments suggest that in some genotypes, SD ratio may vary between leaves and hence it is important to use a defined fully-expanded leaf in analyses of stomatal anatomical patterns.

Growth under dry air conditions led to the formation of smaller stomata with higher densities on both leaf surfaces (Figure 3), whereas SD ratio increased under low RH (Figure 4A), as we and others have found before (Devi and Reddy, 2018; Tulva et al., 2024). Higher SDs should lead to higher maximum gas-exchange potential (Hetherington and Woodward, 2003; Franks and Beerling, 2009; Sack and Buckley, 2016), whereas higher SD ratios may favour increased water loss from leaves due to relatively more stomata localised at the adaxial surface that is exposed to greater evaporative demand (Drake et al., 2019). On the other hand, higher adaxial SDs and SD ratios are common in grasses, including cereals (Muir, 2015; Wall et al., 2022; Samantara et al., 2025), and plants that grow in arid or well-lit habitats (Wood, 1934; Muir, 2018), suggesting that high SD ratios may rather be favourable in drier environments. High SD ratios and SDs do not necessarily mean high water loss and may instead be beneficial under water-limited conditions, allowing rapid gas-exchange during periods when water is available (Mott et al., 1982; Xiong and Flexas, 2020; Ochoa et al., 2024). Smaller stomata developed under dry air conditions may respond faster to changes in environmental conditions (Drake et al., 2013; Durand et al., 2019; Yoshiyama et al., 2024), allowing better control over leaf water loss. Indeed, rice plants with high SDs and smaller stomata were more resilient to drought (Caine et al., 2023). Thus, the formation of smaller denser stomata under dry air conditions may be an adaptive response to combine efficient stomatal responsiveness with high maximum gas-exchange potential.

In addition to SD, stomatal index also increased in the adaxial epidermis under low RH conditions, whereas this effect was present for both mature stomata and total stomatal lineage cells (Figure 4, Supplementary Figure S3), indicating that acclimation to dry air conditions involves adjustment of adaxial stomatal developmental program towards initiation and differentiation of more stomatal lineages and not only reduced cell expansion growth. Contrary to adaxial epidermis, abaxial SI decreased under dry air conditions, as found before in Arabidopsis abaxial epidermis (Tricker et al., 2012), whereas this was only true for mature stomata and not for all stomatal lineage cells (Figure 4). These findings suggest that under low RH, stomatal differentiation into mature functioning stomata is suppressed in the abaxial epidermis, potentially allowing the restriction of stomatal water loss. In line with this, dry air increased the proportion of arrested stomatal lineage cells (Figure 5), suggesting that final differentiation into mature stomata is suppressed under low RH, particularly in the abaxial epidermis. Leaf-side specific changes in SI were also found in cotton, where low RH increased adaxial and did not affect abaxial SI (Devi and Reddy, 2018). In poplar, a negative correlation was found between both adaxial SD and SI, and stomatal response time to dry air (Durand et al., 2019). These results suggest an advantage of stomatal allocation to the adaxial leaf surface under dry air conditions that correlates with faster stomatal closure upon low RH. Potentially, such patterning is triggered via systemic signaling as leaf stomatal patterns are set early in development (Tichá, 1982) and stomatal patterning in newly developing leaves is largely determined by conditions surrounding existing mature leaves (Lake et al., 2001; Coupe et al., 2006). Nevertheless, compared to the relatively large changes in SD under dry air, the increase in adaxial SI, and decrease in abaxial SI were minor (Figures 3-4), indicating that major acclimation of stomatal anatomy under low RH is realised via changes in cell expansion, rather than cell division and differentiation.

Plants grown under dry air conditions were consistently smaller compared with those grown under control conditions and had xeromorphic leaves with high LMAs (Figure 6A, Supplementary Figure S5), despite being well-watered throughout the experiment, as we have found before (Tulva et al., 2023; Tulva et al., 2024). Other studies have found that VPD level may affect plant productivity even more than precipitation (Konings et al., 2017), and high VPD suppresses growth also under well-watered conditions (López et al., 2021), indicating that dry air acts as a stress factor even without its indirect effect through soil moisture. It has been proposed that high VPD suppresses growth partly by restricted A_net_ due to elevated VPD-induced stomatal closure and partly by lowering internal turgor pressure, which inhibits cell division and expansion (reviewed in Novick et al., 2024). We also found a negative relationship between SD and growth (Figure 6B), as detected before in Arabidopsis (Jalakas et al., 2024; Tulva et al., 2024) and rice (Caine et al., 2023), whereas stomatal lineage cell index was also negatively associated with plant size irrespective of growth RH in this study (Figure 6C). The growth reduction associated with excess stomatal density is not explained by higher water loss due to increased stomatal conductance, as the plant lines with more open stomata grew normally, indicating that the smaller size in mutants with high SDs is associated with stomatal abundance *per se*. As stomatal lineage cell index was also negatively associated with plant dry mass, it is likely that stomatal development puts a strain on plant’s resource use even when the effects of leaf expansion on plant size and SD are discounted. Normal stomatal function requires at least one pavement cell between stomata, meaning additional cell divisions in plants with high SIs that are costly for the plant (Bowers, 2018).

In conclusion, we show that acclimation to low air humidity leads to smaller, denser stomata in Arabidopsis. Low RH triggers opposite effects on stomatal development in the adaxial and abaxial leaf surfaces, leading to more stomata being produced on the upper and less on the lower leaf surface. These anatomical changes may prepare the plant for conditions with limited water availability by increasing maximum capacity for gas-exchange via higher stomatal ratios and SDs for when water is available, while allowing fast stomatal closure due to smaller stomata upon water deficit. We also show that pathways that control stomatal development and closure act largely independently and respective alleles can be combined to design desired stomatal traits. As high SDs lead to growth suppression, we conclude that increasing SDs to the levels done here is not helpful as a strategy to improve plant photosynthetic capacity. Whether more moderate increases in SD combined with large stomatal apertures may increase A_net_ and growth under well-watered conditions remains to be tested.

## Materials and methods

### Plant lines used in experiments

The studied *Arabidopsis thaliana* mutant lines were all in Col-0 background (wild-type). Most single mutants were from the European Arabidopsis Stock Center (Scholl et al., 2000): *ghr1-3* (GK_760C07), *ost1-3* (SALK_008068), *slac1-3* (SALK_099139); the *mpk12-4* mutant was from Mikael Brosché (Jakobson et al., 2016), and the *epf1-1epf2-2* double mutant was from Julie Gray (SALK_137549 and GK_673E01, Hunt and Gray, 2009); triple mutants were obtained by crossing the single and double mutants. All plant lines used in experiments are listed in Table 5. Primers used for verifying mutant genotypes are shown in Table S1. Age-matched seed stocks were used for experiments (grown under 50% RH, 23°C; 10:14 h day:night until bolting, then 16:8 h day:night). Plants for all experiments were grown in a 2:1 v/v mixture of peat (OPM 025 W, Kekkilä Oy, Vantaa, Finland) and vermiculite (Vermikuliit Medium 0–4 mm, TopGreen, Vahi, Estonia).

**Table 5.** Plant lines studied, their short names used in this article, and levels of the factors used in ANOVA.

| Plant line | Short name | openness | epf |
| --- | --- | --- | --- |
| Col-0 | Col-0 | wt | 0 |
| *epf1-1 epf2-2* | *epf1/2* | wt | 1 |
| *mpk12-4* | *mpk12-4* | mpk12 | 0 |
| *mpk12-4 epf1-1 epf2-2* | *mpk12-4 epf1/2* | mpk12 | 1 |
| *ghr1-3* | *ghr1-3* | ghr1 | 0 |
| *ghr1-3 epf1-1 epf2-2* | *ghr1-3 epf1/2* | ghr1 | 1 |
| *ost1-3* | *ost1-3* | ost1 | 0 |
| *ost1-3 epf1-1 epf2-2* | *ost1-3 epf1/2* | ost1 | 1 |
| *slac1-3* | *slac1-3* | slac1 | 0 |
| *slac1-3 epf1-1 epf2-2* | *slac1-3 epf1/2* | slac1 | 1 |

### Gas-exchange experiments

Plants for gas-exchange experiments were grown in controlled-environment growth cabinets (Percival AR-22L, Percival Scientific, IA, USA) under standard conditions (10/14 photoperiod with 250 µmol m^-2^ s^-1^ light, relative air humidity 60% during the day and 80% during the night, temperature 23°C during the day and 19°C during the night). For whole-rosette gas-exchange measurements, plants were grown in specialised pots as in Kollist et al. (2007) and watered once a week to ∼60-80% soil water content. Gas-exchange measurements were carried out with a thermostated whole-rosette gas-exchange device designed for Arabidopsis (PlantInvent OÜ, https://www.plantinvent.com/, (Kollist et al., 2007; Hõrak et al., 2017)). Plant gas-exchange was recorded in 23-27 days old plants in measurement cuvettes for 2-3 hours under standard conditions (250 μmol m^-2^ s^-1^ light, ∼55-65% RH, ∼420 ppm CO_2_). Steady-state leaf conductance (g_l_) and net assimilation rate (A_net_) were obtained by averaging respective values over 4 last measured values (average of last 32 minutes in the measurement cuvette). After gas-exchange measurement, leaf 6 was sampled for stomatal density analyses. Pooled data from two independently grown batches of plants is shown in figures.

For leaf gas-exchange measurements, plants were grown in 110 ml pots and watered to soil water holding capacity ∼ 2 times a week. Adaxial and abaxial stomatal conductance (g_s_) measurements were carried out with a leaf porometer (LI-600, LI-COR Environmental) on one leaf each of 4-5 weeks old plants under standard conditions. Stomatal conductance ratio (g_s_ ratio) was calculated as the ratio of adaxial and abaxial stomatal conductances. The same leaf that was analysed by porometry was sampled for stomatal density analyses. Pooled data from two independently grown batches of plants is shown in figures.

### Growth experiment

Growth experiment was carried out similar to Tulva et al. (2024). Plants were grown in 50x29 cm trays filled with 6 cm deep soil. Water was added to the outer tray and absorbed by the soil in the inner tray through 40 holes drilled into the inner tray’s bottom. Soil in each tray was covered with a sheet of mulch cloth (Baltic Agro AS, Lehmja, Estonia). The cloth had ten 5 mm holes 10 cm apart, into which plants, one plant of each studied line, were sown in a pattern randomized for each tray (Figure S6).

Soil was weighed and sampled from each tray during filling; the soil sample was then dried for a week at 70°C to determine soil water content (SWC). 3 l of water was immediately added to each tray, taking SWC to ∼85% for sowing. After sowing, the trays were wrapped in cling film which was removed one week later. Trays were moved to two Snijders Microclima Arabidopsis MCA1600-3LP6-E growth cabinets (Snijders Scientific, Tilburg, Netherlands) set to maintain air temperature 23°C, 19°C at night, RH 60% at daytime 80% at night (VPD 1.1 kPa vs 0.4 kPa), with 10:14h day:night cycle. Light intensity in the cabinets was 250 μmol m^-2^ s^-1^. Starting at the second week after sowing, each tray was weighed daily to calculate soil water content.

Whenever SWC in any tray was below 70%, water was added to the outer tray to take SWC to 75%. Trays were shuffled between shelves, positions on a shelf, and growth cabinets daily.

Fourteen days after sowing, one of the growth cabinets was switched to “dry” regime, which was 40% RH at daytime and 50% at night (VPD 1.7 kPa vs 1.1 kPa), otherwise identical to the control. After that, the trays were still shuffled around within their cabinets throughout the experiment, but no more between. Daily weighing of the trays continued with the above algorithm, ensuring a long-term equal SWC in all trays.

On the 39th day, aboveground parts of all plants were harvested. No plants showed signs of bolting yet. The 12th leaf was separated, photographed for its area, and weighed for fresh mass, then used for epidermal imaging as described below. The rest of the rosette was weighed for its fresh mass. Finally, the rosettes were dried in an oven until stable mass (4-5 days) and weighed again. Plant dry mass was obtained from the rosette dry mass, corrected for the missing 12th leaf by scaling with the ratio of respective fresh masses. Leaf mass per area (LMA) was obtained as 12th leaf dry mass/12th leaf area.

The growth experiment was conducted twice, with the growth cabinets swapped. Each run consisted of 24 trays, twelve in each treatment. A few plants and, in one run, two whole trays were lost to handling accidents, resulting in a total of 20-23 plants per line per treatment.

### Epidermal patterning analyses

Sampled leaves were cut in half along the midvein and epidermal impressions were generated from the adaxial surface of one half and from the abaxial surface of the other via dental silicone (Speedex light body, Coltene/Whaledent AG for adaxial and oranwash L, Zhermack for abaxial) as described before (Jalakas et al., 2024). Nail varnish was applied to the dental silicone, allowed to dry and transferred to microscope slides via transparent tape. A video encompassing different focal plains was captured from an area of ∼0.28 mm^2^ in the central part of each impression (with respect to leaf tip and base, and central vein and edge) with Kern OBF 133 microscope and ODC 832 camera (Kern & Sohn GmbH). Analyses of cell density and size were carried out from the obtained videos with ImageJ (Schneider et al., 2012). Stomatal density (SD) was defined as the number of mature stomata per mm^2^; total stomatal density (total SD) was defined as the sum of adaxial and abaxial SDs; stomatal lineage cell density (LCD) was defined as the number of all stomatal lineage cells (arrested stomatal precursors + mature stomata) per mm^2^; pavement cell density (PCD) was defined as the number of pavement cells per mm^2^; stomatal index (SI) was defined as the proportion of mature stomata from all epidermal cells (SD/(LCD+PCD)); stomatal lineage cell index (LCI) was defined as the proportion of stomatal lineage cells from all epidermal cells (LCD/(LCD+PCD)); stomatal ratio was defined as the ratio of adaxial and abaxial SDs; stomatal lineage cells ratio (LCD ratio) was defined as the ratio of adaxial and abaxial LCDs.

### Statistical analyses

Genotype data was split into two factors, a binary variable “epf” which encoded presence of lack of *epf1/2* double mutation, and a five-level factor “openness” that took one of values “wt”, “mpk12”, ”ost1”, ”slac1”, or “ghr1” that coded any other mutation involved. In the growth experiment data, growth conditions were encoded as one more binary factor “RH”, taking values “control” or “dry”.

Statistical analyses were carried out with R version 4.2.1 (r-project.org). With variables that passed Levene’s test for homogeneity of variance, analysis of variance with type III sums of squares was used by employing function Anova(…,type=”III”) from the library car (https://r-forge.r-project.org/projects/car/) and contrasts were explored with Tukey’s HSD test for unbalanced sample (library car, https://myaseen208.github.io/agricolae/). Where Levene’s test failed (p<0.05), logarithm of the variable was analogously tested for homogeneity of variance, and used in ANOVA if passed. This, too, failing, Kruskal-Wallis rank test was used instead, with the ten-level genotype as predictor and Dunn’s test for post-hoc.

### Accession numbers

AGI accession numbers for genes studied in this article are AT2G20875 (EPF1), AT1G34245 (EPF2), AT2G46070 (MPK12), AT4G20940 (GHR1), AT4G33950 (OST1), and AT1G12480 (SLAC1).

## Supplementary data

Supplementary table S1. Individual mutant lines used in this study and primers used for genotyping them.

Supplementary table S2. Power regressions for SD vs SL relationship.

Supplementary table S3. Tukey test of the interaction of growth RH and stomatal openness mutations on adaxial stomatal length.

Supplementary table S4. Tukey test of the openness mutations on LMA.

**Supplementary Figure S1.** Nonsignificant relationship between A_net_ and g_l_ or total SD in whole-rosette gas-exchange experiments.

**Supplementary Figure S2.** Proportion of adaxial g_s_ and SDs in leaf gas-exchange experiments.

**Supplementary Figure S3.** Total stomatal lineage cell ratio and stomatal lineage cell indices in growth experiments.

**Supplementary Figure S4.** Representative epidermal images of fully expanded leaf 12 from growth experiments.

**Supplementary Figure S5.** LMA in the growth experiments.

**Supplementary Figure S6.** Representative images of trays before treatment start and at the end of the experiment under normal and low RH.

## Acknowledgements

We thank Julie Gray for the *epf1epf2* double mutant seeds and Luke Fountain for help with genotyping F1 plants. We thank Mikk Välbe, Sára Babincová, and Madli Johanna Veigel for help with experiment maintenance and harvesting, and Ebe Merilo for scientific discussions and comments on the manuscript.

## Author contributions

I.T., P.J., and H.H. designed the study, I.T., P.J., E.I., A.W., P.K.O., and H.H. performed experiments and analyzed data, I.T. and H.H. wrote the manuscript, all authors commented, edited and approved the final manuscript.

## Data availability

All analysed data described in this manuscript is presented in main and supplementary figures and tables. Raw data will be submitted to a data repository upon acceptance of the manuscript.

