## Supplementary Figures for "Low relative air humidity leads to smaller, denser stomata and higher stomatal ratios in Arabidopsis"

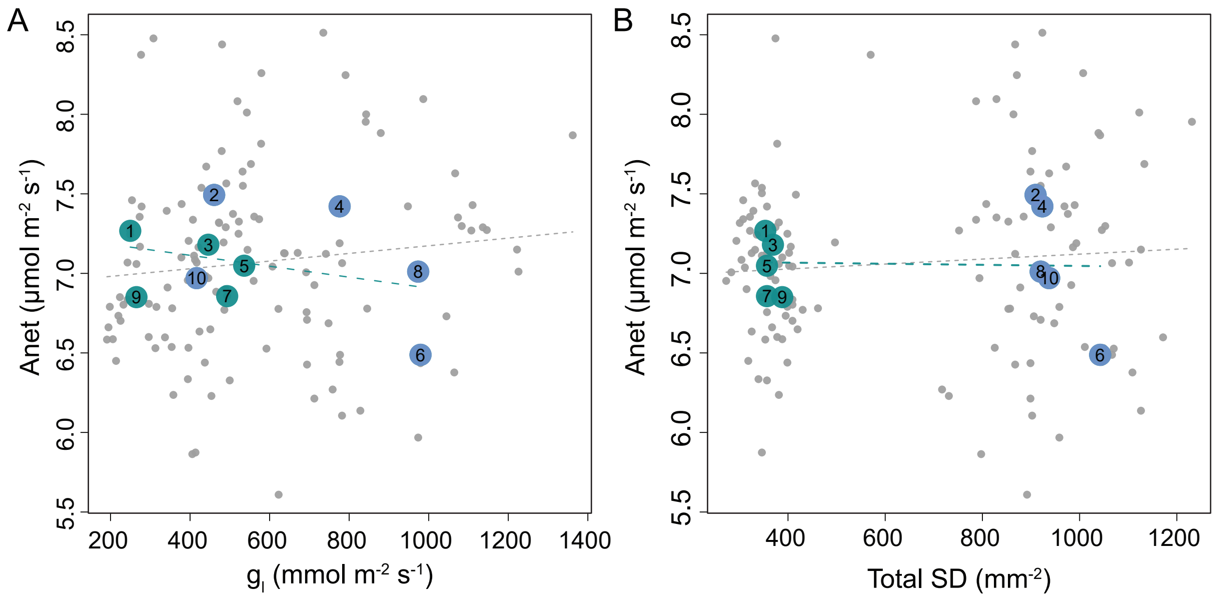


**Supplementary Figure S1.** Relationship between net photosynthesis and stomatal properties (whole-plant leaf conductance in (A), the sum of abaxial and adaxial stomatal densities in (B)) in the gas exchange experiment. Gray points are individual plants, numbered circles denote each genotype's medians. Slopes of the linear relationships, illustrated with hatched lines, did not differ significantly from zero.

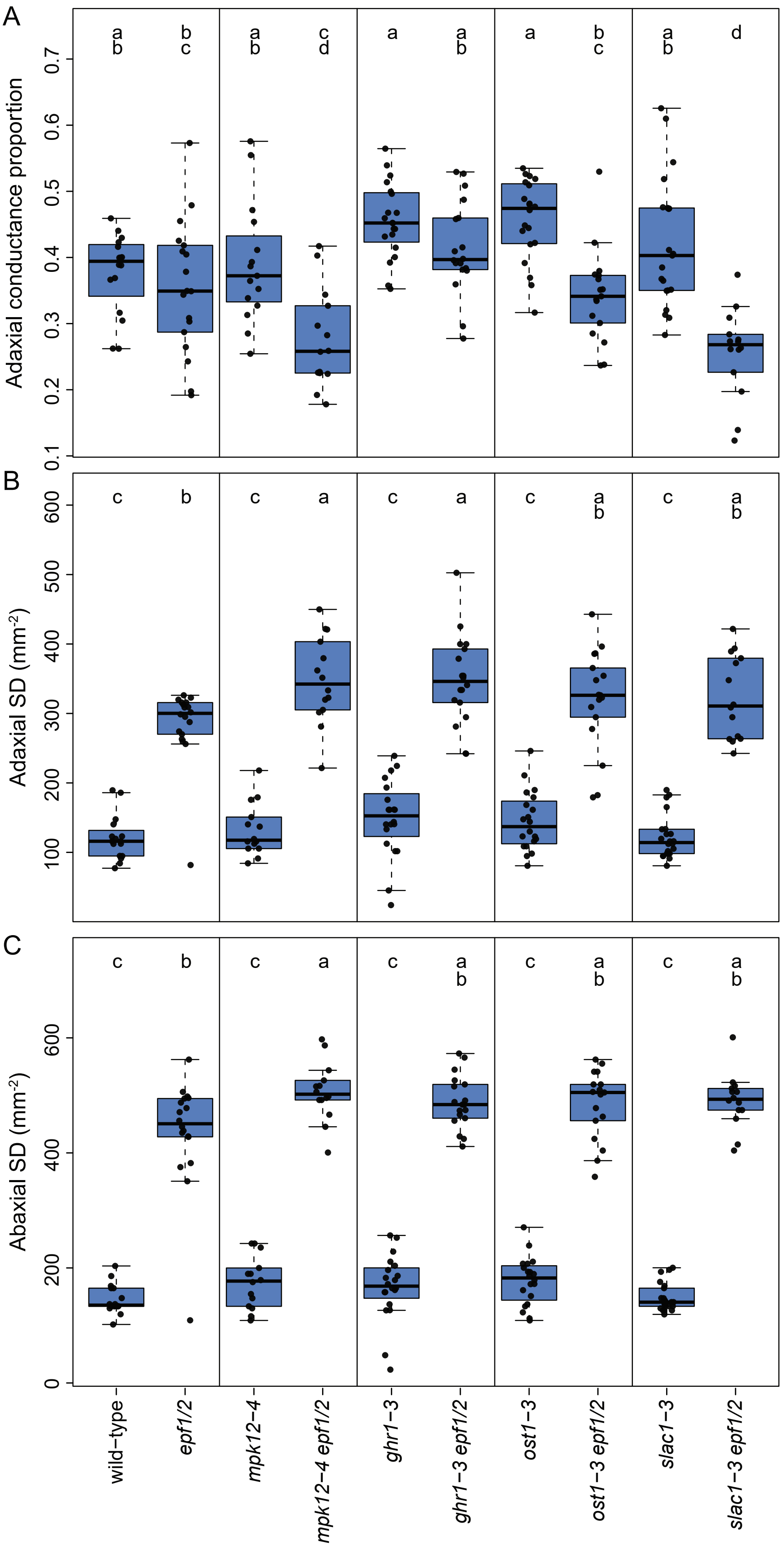


**Supplementary Figure S2.** Proportion of adaxial conductance in the whole leaf conductance (A) in the porometry experiment, and stomatal densities at the adaxial and abaxial leaf surfaces (B)-(C). For each plant line, the box denotes the lower to upper quartile range, with the median marked with a line. The whiskers span the non-outlier range, and black dots mark individual data points (n=14-22). Letters above the boxes denote Tukey post-hoc test results, with plant lines not sharing a letter differing from each other at $p<0.05$.

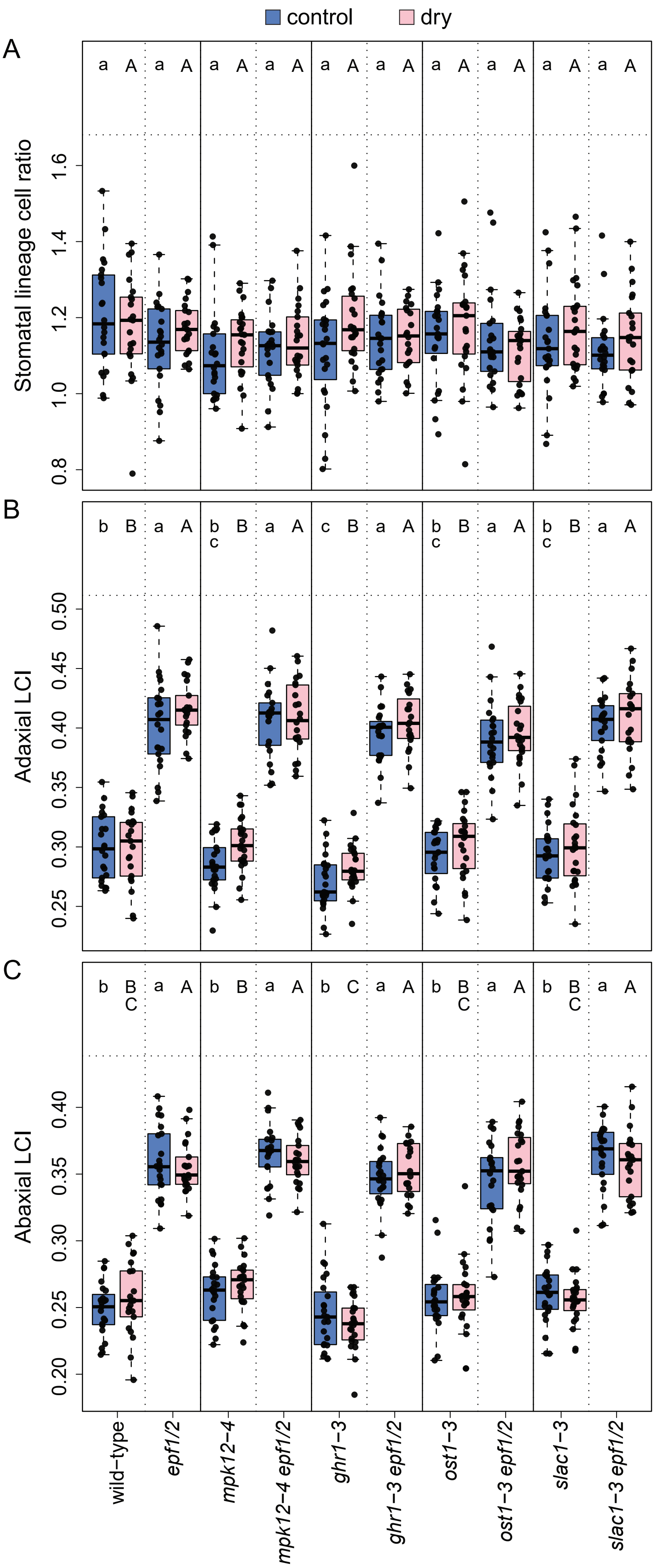


**Supplementary Figure S3.** Stomatal lineage cells (i.e. mature stomata plus stomatal precursors) in the growth experiment. (A), ratio of stomatal lineage cell abundance between adaxial and abaxial leaf surfaces. (B)-(C), proportion of stomatal lineage cells from all epidermal cells (LCI) at the adaxial and abaxial leaf surface. The boxes denote the lower to upper quartile range, with the median marked with a line. The whiskers span the non-outlier range, and black dots mark individual data points (n=20-23). Small letters above the blue control boxes denote Tukey post-hoc test results, with plant lines not sharing a letter differing from each other at $p<0.05$ (one-way ANOVA for genotype differences involving only control plants), and capital letters analogously for the low RH treatment.

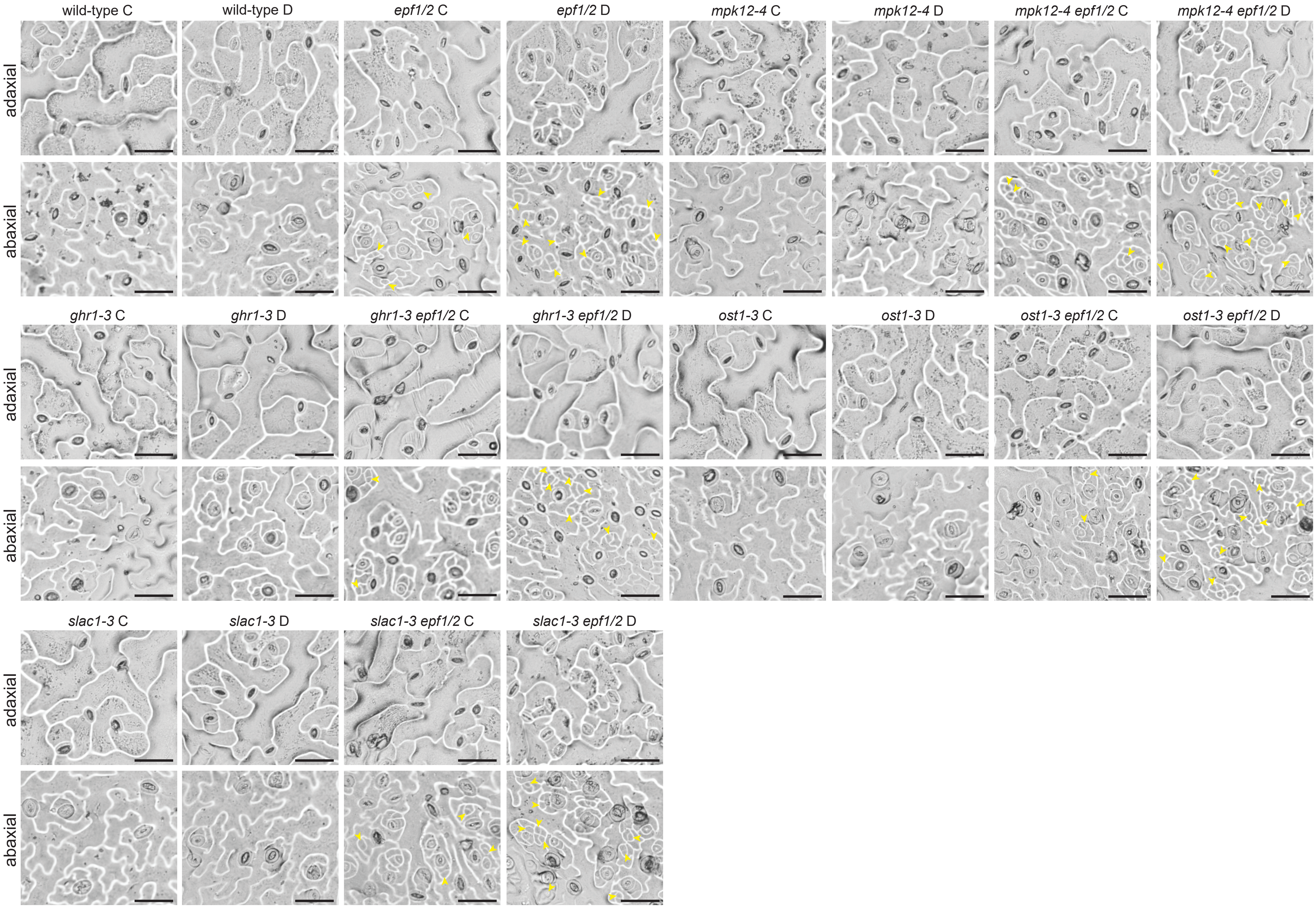


**Supplementary Figure S4.** Representative epidermal images of fully expanded leaf 12 from growth experiment (normal (C) and low RH (D)). Yellow arrowheads indicate precursor cells. The scale bar represents 50 μm.

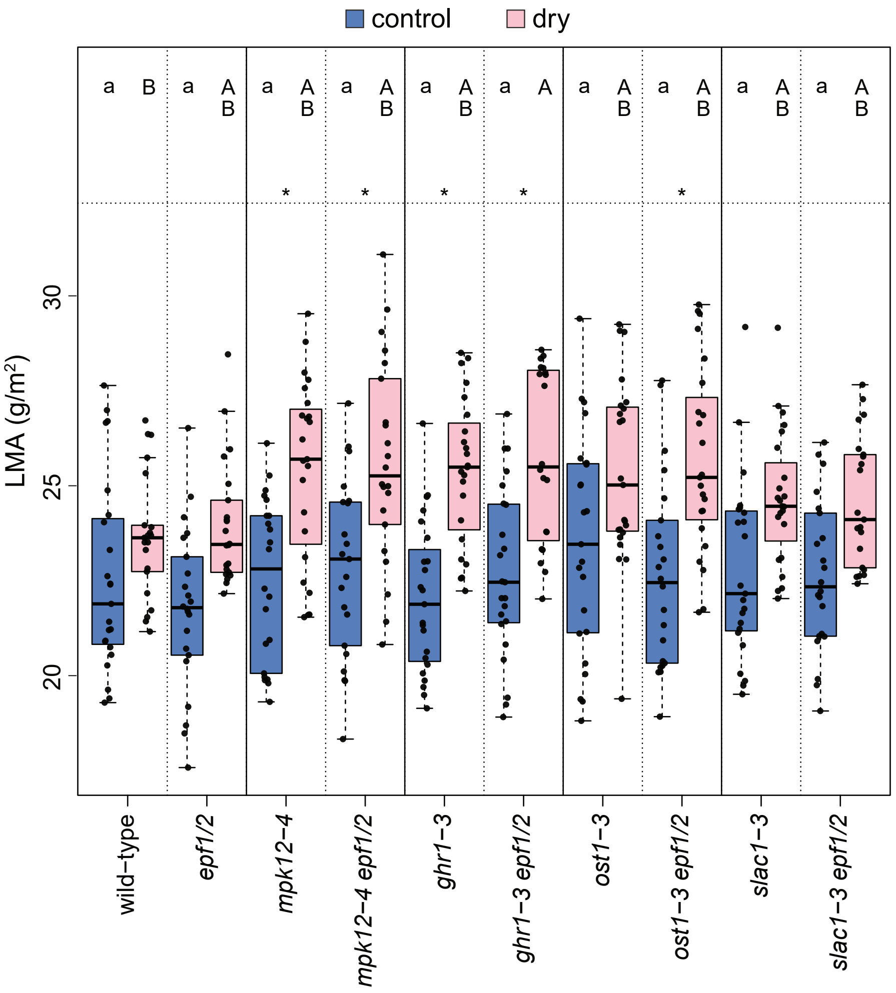


**Supplementary Figure S5.** Leaf mass per area in the growth experiment. The boxes denote the lower to upper quartile range, with the median marked with a line. The whiskers span the non-outlier range, and black dots mark individual data points (n=20-23). Asterisk above a pair of boxes means significant pairwise difference between plants grown in low RH and control conditions according to Tukey test (factorial ANOVA involving all treatment and line combinations). Small letters above the blue control boxes denote Tukey post-hoc test results, with plant lines not sharing a letter differing from each other at $p<0.05$ (one-way ANOVA for genotype differences involving only control plants), and capital letters analogously for the low RH treatment.

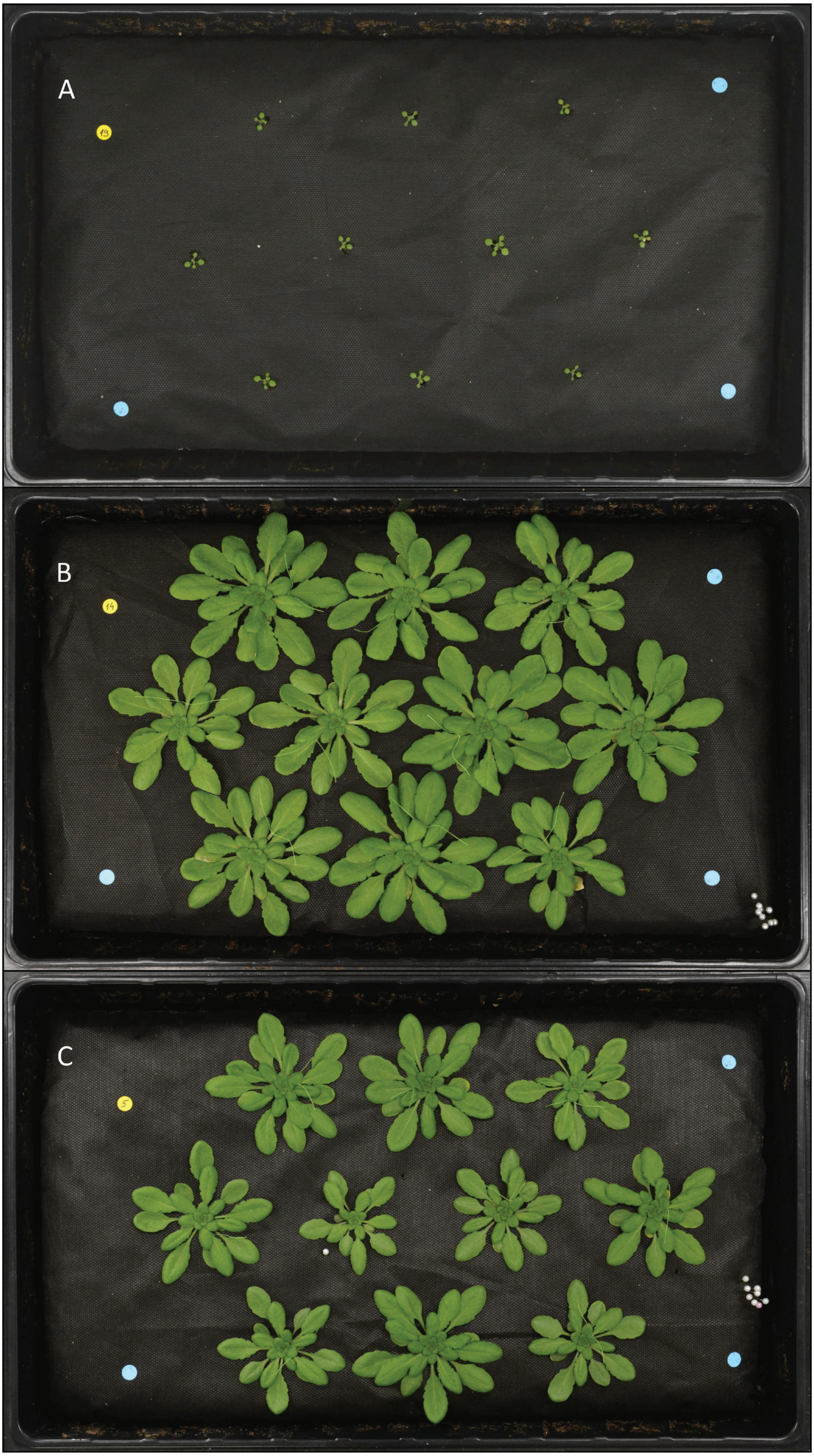


**Supplementary Figure S6.** Example photos of the growth experiment trays on the 14th day after sowing (when differential treatment started, A) and the day before harvesting: (B), control; (C), low RH. The threads had been attached ~a week prior to mark the 12th leaf of each plant that would be analyzed for stomatal anatomy.
