## Supplementary Tables for "Low relative air humidity leads to smaller, denser stomata and higher stomatal ratios in Arabidopsis"

**Supplementary table S1.** Individual mutant lines used in this study and primers used for genotyping them.

| Mutant line | Line name | T-DNA border primer | Left primer (LP) | Right primer (RP) | Genotyping principle |
|---|---|---|---|---|---|
| *epf1-1* | SALK_137549 | ATTTTGCCGATTTCGGAAC | GGTGCATGTTCGACACTCTTC | CATGGTCATGTCCCGGAGAAGC | Product with LP + RP from wild-type allele, and with T-DNA border primer + RP from mutant allele |
| *epf2-2* | GABI_673E01 | ATATTGACCATCATACTCATTGC | ATGACGAAGTTTGTACGCAAG | AATCTGATTCGGTCGTGTC | Product with LP + RP from wild-type allele, and with T-DNA border primer + RP from mutant allele |
| *mpk12-4* | WT allele | not a T-DNA mutant | GCTAGATCTATCAAGTGTTCCTG | GGTTAGGATCAAAGACGAGC | See Jakobson et al. 2016 |
| | Mutant allele | | CATCATCACTCGAGAGAATTTTCG | CTCCGGTCATCGCCGTTATC | |
| *ghr1-3* | GK_760C07 | ATATTGACCATCATACTCATTGC | GAGCTTAGAGAGACAATGTCATTC | CTCCTTATCTGGACCTTTGCC | Product with LP + RP from wild-type allele, and with T-DNA border primer + RP from mutant allele |
| *ost1-3* | SALK_008068 | ATTTTGCCGATTTCGGAAC | CATATCTTTAGACGAGGGGCC | GTGAGTGGTCCAATGGATTTG | Product with LP + RP from wild-type allele, and with T-DNA border primer + RP from mutant allele |
| *slac1-3* | SALK_099139 | ATTTTGCCGATTTCGGAAC | AACTTCTTCTTCGCTCCTTGG | GACCATTTCTTTGCCTGTTTG | Product with LP + RP from wild-type allele, and with T-DNA border primer + RP from mutant allele |

**Supplementary table S2.** Power regressions shown in Figure 3 (E)-(F). SL, stomatal length; SD, stomatal density. Intercept and slope are for the linear regression between ln(SD) and ln(SL).

| Leaf side | EPF state | Intercept ± SE | Slope ± SE | Equation |
|---|---|---|---|---|
| adaxial | wt | 3.852 ± 0.068 | -0.157 ± 0.014 | $SL = 47.1 * SD^{-0.157}$ |
| | epf1/2 | 3.933 ± 0.095 | -0.142 ± 0.016 | $SL = 51.1 * SD^{-0.142}$ |
| abaxial | wt | 3.940 ± 0.085 | -0.157 ± 0.017 | $SL = 51.4 * SD^{-0.157}$ |
| | epf1/2 | 4.187 ± 0.189 | -0.164 ± 0.030 | $SL = 65.8 * SD^{-0.164}$ |

**Supplementary table S3**. Tukey test of the interaction of growth RH and stomatal openness mutations on adaxial stomatal length (Table 3). Shown are mean and its standard error for each stomatal openness and treatment combination, with EPF levels pooled. Different letters denote statistically significant differences.

| Openness, treatment | Mean±SE |
|---|---|
| wt, control | 22.66 ± 0.20 c |
| wt, dry | 21.50 ± 0.17 d |
| mpk12, control | 23.87 ± 0.18 ab |
| mpk12, dry | 21.69 ± 0.15 d |
| ghr1, control | 23.18 ± 0.26 bc |
| ghr1, dry | 20.45 ± 0.16 e |
| ost1, control | 24.29 ± 0.15 a |
| ost1, dry | 21.84 ± 0.16 d |
| slac1, control | 23.67 ± 0.20 ab |
| slac1, dry | 21.16 ± 0.17 de |

**Supplementary table S4**. Tukey test of the openness mutations on LMA (Table 3). Shown are mean and its standard error for each stomatal openness, with EPF levels and treatments pooled. Different letters denote statistically significant differences.

| Openness | Mean±SE |
|---|---|
| wt | 22.98 ± 0.23 b |
| mpk12 | 24.13 ± 0.30 a |
| ghr1 | 23.93 ± 0.28 a |
| ost1 | 24.30 ± 0.30 a |
| slac1 | 23.66 ± 0.23 ab |
